# A Local Measure of Symmetry and Orientation for Individual Spikes of Grid Cells

**DOI:** 10.1101/425637

**Authors:** Simon N. Weber, Henning Sprekeler

## Abstract

Grid cells have attracted broad attention because of their highly symmetric hexagonal firing patterns. Recently, research has shifted its focus from the global symmetry of grid cell activity to local distortions both in space and time, such as drifts in orientation, local defects of the hexagonal symmetry, and the decay and reappearance of grid patterns after changes in lighting condition. Here, we introduce a method that allows to visualize and quantify such local distortions, by assigning both a local grid score and a local orientation to each individual spike of a neuronal recording. The score is inspired by a standard measure from crystallography, which has been introduced to quantify local order in crystals. By averaging over spikes recorded within arbitrary regions or time periods, we can quantify local variations in symmetry and orientation of firing patterns in both space and time.

## 1 INTRODUCTION

Neurons in the entorhinal cortex are often classified based on their spatial firing patterns. Grid cells, in particular, have attracted a lot of attention, because of their highly symmetric hexagonal activity (Fyhn et al., 2004; Hafting et al., 2005). To decide whether a given cell should be classified as a grid cell or a non-grid cell (Diehl et al., 2017; Hardcastle et al., 2017; Krupic, Burgess, and O’Keefe, 2012), researchers need a grid score: a number that quantifies the degree of hexagonal spatial symmetry in the firing pattern of the cell. Since the first reports of grid cells, the community has developed and refined a standard grid score (Barry and Burgess, 2017; Langston et al., 2010; Sargolini et al., 2006a; Wills et al., 2010), which relies on the following procedure. Spike locations are transformed into a rate map. The peaks in the spatial autocorrelogram of this rate map are used to obtain measures of the grid spacing and the grid orientation. The auto-correlogram of the rate map is then cropped, rotated and correlated with its unrotated copy. A grid score is finally obtained from the resulting correlation-vs-angle function at selected angles (for details see Section S.3). This procedure assumes a consistent hexagonal pattern throughout the arena and results in a single grid score for the entire firing pattern.

While this grid score has many benefits, it is inherently global and ignores local properties of firing patterns. Recent studies have shown, however, that the firing patterns of grid cells show interesting local effects, such as drifts in orientation (Stensola et al., 2015) or local defects of the hexagonal symmetry (Krupic, Bauza, Burton, Barry, et al., 2015; Krupic, Bauza, Burton, and O’Keefe, 2018). These results call for new methods for quantifying the *local symmetry* and the *local orientation* of grids.

Here, we introduce a method that assigns both a grid score and an orientation to *each individual spike* of a neuronal recording. This spike-based score is inspired by a standard measure from crystallography (Halperin and Nelson, 1978), which has been introduced to quantify local order in crystals. By averaging over spikes recorded within arbitrary regions or time periods, we can quantify local variations in symmetry and orientation of firing patterns in both space and time. A high spatial and temporal resolution can be achieved by averaging over multiple cells.

To illustrate the potential of our method, we apply it to both artificially generated data and neuronal recordings from grid cells. In the following, we first introduce the method in detail. Comparing it to the conventional correlogram-based grid score (Langston et al., 2010), we show that both methods perform similarly when quantifying and classifying grid properties globally. We then illustrate the benefits of our spike-based score to analyze and visualize local properties of spike patterns in space and time.

## 2 METHODS

The suggested grid score relies on a measure that is used to describe symmetry in crystals (Halperin and Nelson, 1978). Neuronal recordings differ from crystalline arrangements of atoms in several respects and therefore call for some modifications. Firstly, crystals form a symmetric arrangement of neighboring atoms, whereas grid cells show a symmetric arrangement not of spike locations, but of firing fields, each of which corresponds to many spikes. This requires a modified definition of neighborhood. Secondly, crystals consist of many atoms, which show rare deviations from a symmetric arrangement. In contrast, grid cells show relatively few firing fields in an often highly non-symmetric arrangement. This requires a modification to avoid false positives arising from other symmetries. In the following, we first describe the basic idea of the score and then introduce the modifications that are necessary to adjust it to neuronal recordings.

*Python* code of the method is available from our repository https://gitlab.tubit.tu-berlin.de/simonweber/gridscore (Weber, 2018) under the GNU General Public License v3.0. We also encourage readers to try the suggested grid score on their own data on our website: http://gridscore.cognition.tu-berlin.de/.

### 2.1 Measuring M-fold symmetry in crystals

The angular symmetry of an arrangement of N neighbors around a given atom in a crystal can be quantified by a complex number (Halperin and Nelson, 1978):

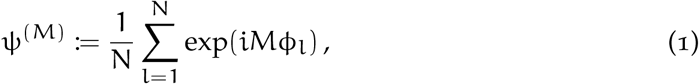

where *i* is the imaginary unit and ϕ_l_ is the angle between an arbitrary reference vector and the vector pointing from the atom in question to its l-th neighbor (Halperin and Nelson, 1978). In the following, we will use the horizontal axis from left to right as the reference vector (Figure 1a). M is a natural number that characterizes the symmetry of interest. For instance, M = 6 corresponds to a hexagonal arrangement of atoms and M = 4 to a quadratic arrangement. The computation of this score can be visualized as an addition of vectors in the complex plane (Figure 1a).

**Figure 1:**
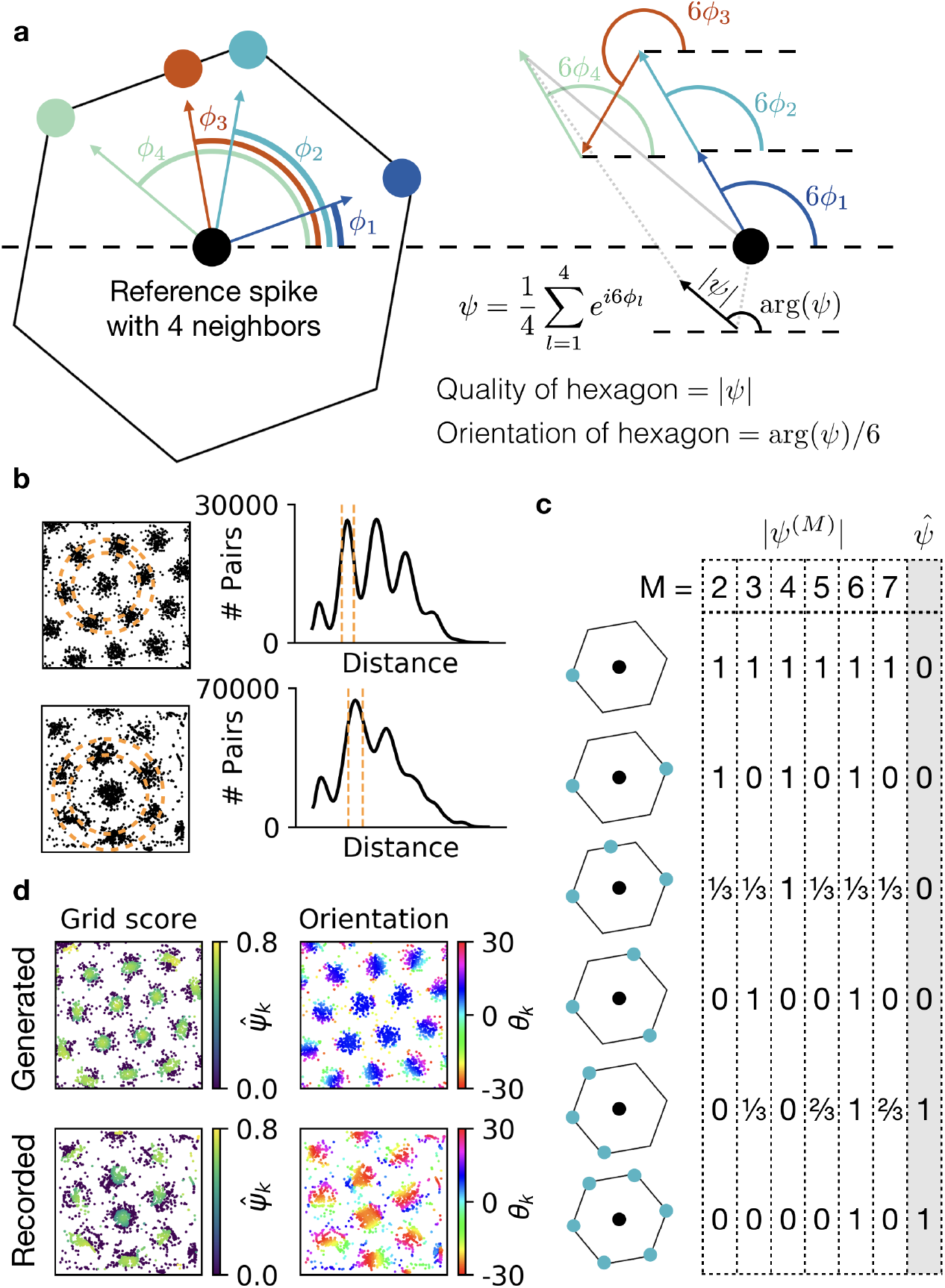
a) Illustration of the ψ measure for 6-fold symmetry. Left: A reference ‘atom’ (black disc) with four neighbors (colored discs). Arcs show the angles, ϕ_l_, to a reference axis (dashed black line) for all neighbors l ∈ {1, 2, 3, 4}. The blueish neighbors lie on the corners of a hexagon. The red neighbor is a defect to this hexagonal structure. Right: Sum of four unit vectors with directions given by 6ϕ_l_ (colored arcs). Vectors associated with neighbors that lie on the corners of the same hexagon (blueish colors) point in the same direction. The vector associated with the defect (red) points in a different direction. The length |ψ| of the resulting vector, normalized by the number of neighbors (black arrow), quantifies how much the reference atom is embedded in a hexagonal structure. The direction arg(ψ) of the vector—divided by 6 to reverse the previous rotation—indicates the orientation of the hexagon. **b)** Detection of the neighborhood shell. Locations of spikes of a grid cell and smoothed histogram of the pairwise distances between all spikes. Dashed lines indicate the automatically detected neighborhood shell. Top: Generated data. Bottom: Experimental recording kindly provided by Stensola et al. (2015). **c)** Preventing false positives by discarding other symmetries. Example |ψ^(M)^| and ψ̂ values for a spike (black) with few neighbors (blue) in six different configurations. **d)** Same spike maps as in b, color-coded with the spike-based grid score ψ̂_k_ (left) and the spike-based orientation 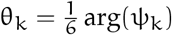 (right).

The absolute value |ψ^(M)^| ranges from 0 to 1, and larger values indicate a more pronounced M-fold symmetry. If the N neighboring atoms lie on the corners of a regular M-polygon, the angle 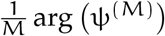 is the orientation of this polygon (Figure 1a). The function arg(x) maps to values between −π and π radians, so the resulting orientation is between −π/M and π/M. For hexagonal symmetry, this corresponds to values between −30 and 30 degrees.

### 2.2 Defining the neighborhood of a spike

The ψ measure has been introduced to quantify symmetry in crystalline structures where *neighborhood* is clearly defined. For example, the nearest neighbors of a reference atom are all atoms within a distance of one lattice constant. For a given spike in a grid pattern, however, the nearest neighbors are most likely other spikes from the same grid field. To assess whether a given spike is part of a symmetric grid, we are hence interested not in neighboring spikes, but in spikes in neighboring grid fields, i.e., spikes that are roughly one grid spacing away from the spike in question. To identify these relevant spikes, we define a *neighborhood shell*—a two dimensional annulus (Figure 1b). We then center this neighborhood shell at the location of the spike in question and consider all spikes within the shell as its neighbors. Assuming that the grid spacing does not change drastically across the environment, we use the same shell size for all spikes.

To identify a suitable size of the shell, we use the histogram of pairwise distances between all spike locations. For periodic firing patterns, this histogram has pronounced peaks (Figure 1b). The first peak characterizes the size of the grid fields. The second peak corresponds to the grid spacing ℓ, and further peaks arise at multiples of the grid spacing. We therefore set the inner radius of the neighborhood shell to 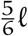 and the outer radius to 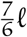 (Figure 1b). To reduce the noise arising from finite numbers of spikes, we determine the peaks from a smoothed histogram, obtained by applying a Gaussian filter with a standard deviation of 1% of the maximal pairwise distance. The suggested shell width of 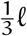 is motivated by the typical size of grid fields, but the suggested grid score is robust to changes of the shell width. Scaling the shell width with the grid spacing is commensurate with the experimental finding that the ratio of grid spacing and grid field size is constant (Brun et al., 2008).

The detection of the correct neighborhood shell as the second peak of the histogram of pairwise distances is a crucial step in the computation of the grid score. For noisy grid patterns, the first peak of the histogram—which indicates the size of the grid fields—is sometimes hard to detect, leading to a potential misclassification of the second peak as the first, and consequently a wrong neighborhood shell (Figure 3f). In data sets where this is the case (Figure 6, Figure 4a,b,c,d), we use the following remedy: We define a rough cutoff distance that is larger than the grid field size and smaller than the grid spacing, and use the first peak above the cutoff to determine the shell size. For the analyzed grid patterns, taking 15% of the side length of the arena for this cutoff worked reliably.

In the dataset with very noisy and elliptic grids (Figure 4e,f,g,h,i,j), peak detection can fail for many cells. In this case, we exploit the fact that nearby grid cells have a similar grid spacing (Hafting et al., 2005) and determine the neighborhood shell from less noisy grid cells of the same population.

### 2.3 Comparing to other symmetries prevents false positives

A large 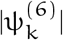 value indicates that the neighbors of the k-th spike are arranged on the corners of a hexagon, but it does not require that all corners of the hexagon are occupied. If, for example, the firing fields of a cell are arranged on a line, this leads to high |ψ^(6)^| values although the structure does not resemble a grid (Figure 1c). This can lead to a wrongful classification of a cell as a grid cell, i.e., to false positives. To avoid high local grid scores without an actual 6-fold symmetry, we compute 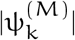 for a range of symmetries (M ∈ {2, …, 7}) and only assign a non-zero grid score to spike k if the hexagonal symmetry is the strongest:

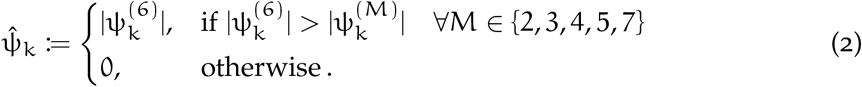

We refer to 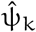 as the *grid score of spike* k. Comparing to symmetries with M > 7 did not have an effect on our results, so we restricted M to a maximum of 7.

A global grid score ψ and the global grid orientation Θ for a given cell is calculated by averaging over the respective properties of the individual spikes. A summary of the algorithm is shown in Algorithm 1.

### 2.4 Tools to evaluate the spike-based grid score

In the following, we compare the suggested local grid score to a conventional correlation-based grid score. Details on the calculation of this score, on the shuffling procedure for grid cell classification, and on the generation of artificial spike data can be found in Section 4.7.

#### Algorithm 1: Algorithm to compute spike-based grid scores 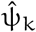 and orientations θ_k_ and their averages ψ and Θ.

**Input:** Two dimensional spike locations **x**_k_, where k ∈{1, …, N^spikes^}

// Get the central radius of the neighborhood shell

Compute the smoothed histogram of pairwise distances between all spike locations and determine the position ℓ of the second peak of this histogram.

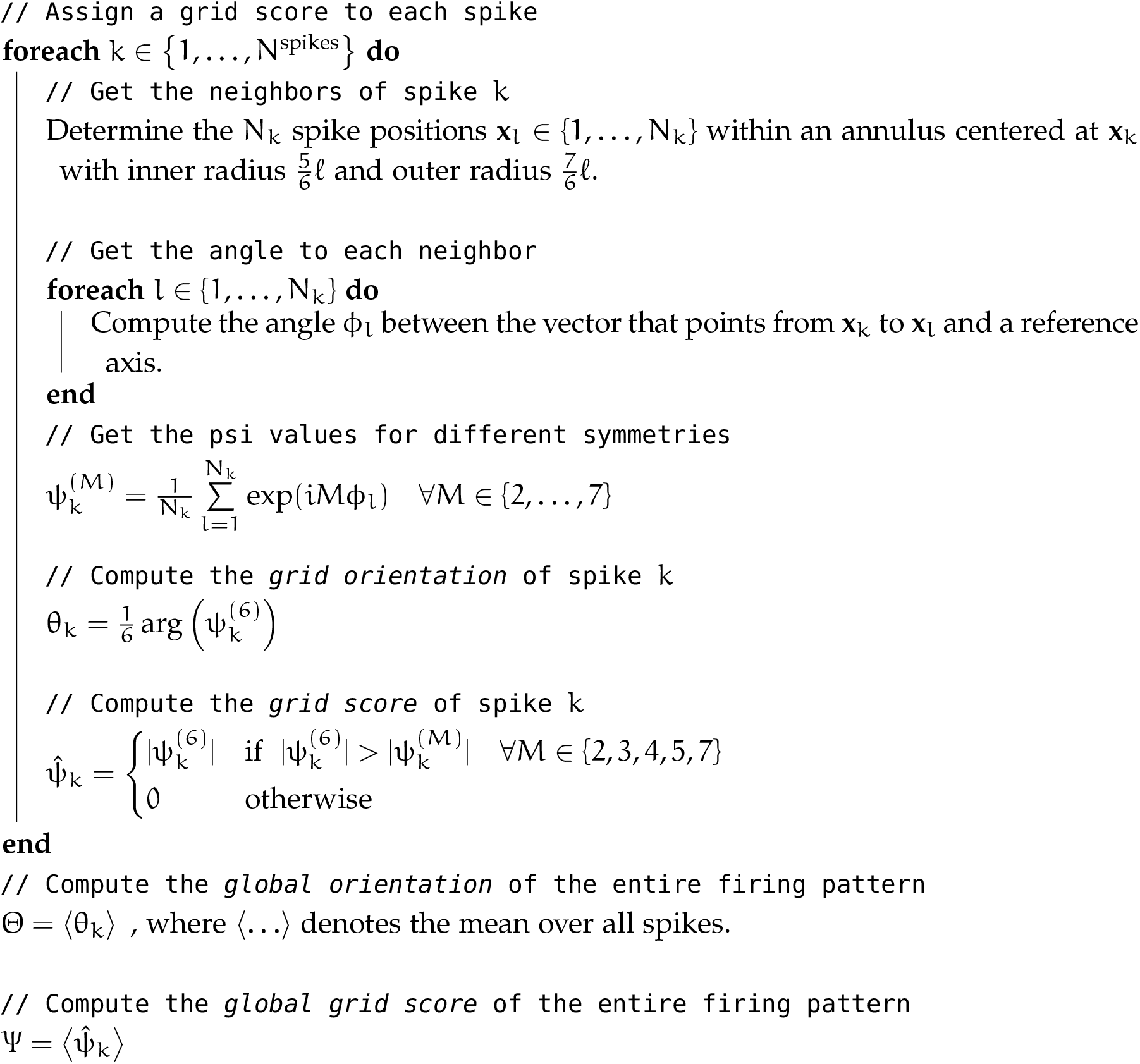

## 3 RESULTS

To evaluate the spike-based grid score, we apply it to generated and experimentally recorded firing patterns. We begin by contrasting the mean grid score of all spikes, ψ (see Algorithm 1), and the correlogram-based grid score, ρ (see Section S.3), and show that they are similarly powerful for quantifying the symmetry of grid patterns and classifying grid cells. We then demonstrate how the individual spike scores, 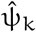 (see Algorithm 1), can be used to systematically analyze local distortions of gridness and orientation in space and time.

### 3.1 The mean of the spike-based score characterizes global grid properties

Grid scores are typically used for two purposes: Quantifying the hexagonality, i.e., the *gridness*, of the spike map and classifying a cell as a grid cell. Given that the patterns of grid cells are rarely perfectly hexagonal, a grid score should classify a cell as a grid cell even if the hexagonal pattern is weakly distorted. For example, a grid score needs to be robust against small shifts of grid field locations or elliptic distortions like shearing. At the same time, a grid score should quantify such distortions by returning lower values for stronger distortions.

#### Quantifying gridness

To study the range of values for the ρ score and the score, we generate spike locations with different degrees of perturbation by randomly shifting the locations of grid fields arranged on a perfect hexagonal grid (Figure 2a, b, see Section S.2 for details). The less hexagonal the arrangement of firing fields, the fewer spikes have a large 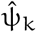 score (Figure 2a). The ψ score, i.e., the mean over all spikes, and the correlogram-based ρ score, both decay with increasing noise on field locations (Figure 2c, results from 100 random realizations of spike patterns for each noise level). ψ scores are around 0.35 for perfect grids and decrease to zero for random field locations. ρ scores are around 1.5 for perfect grids and decrease to −0.5 for random field locations. ψ scores close to 1 are never reached, because many spikes at the edges of grid fields have an individual grid score of 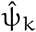 = 0 (Figure 2a).

**Figure 2:**
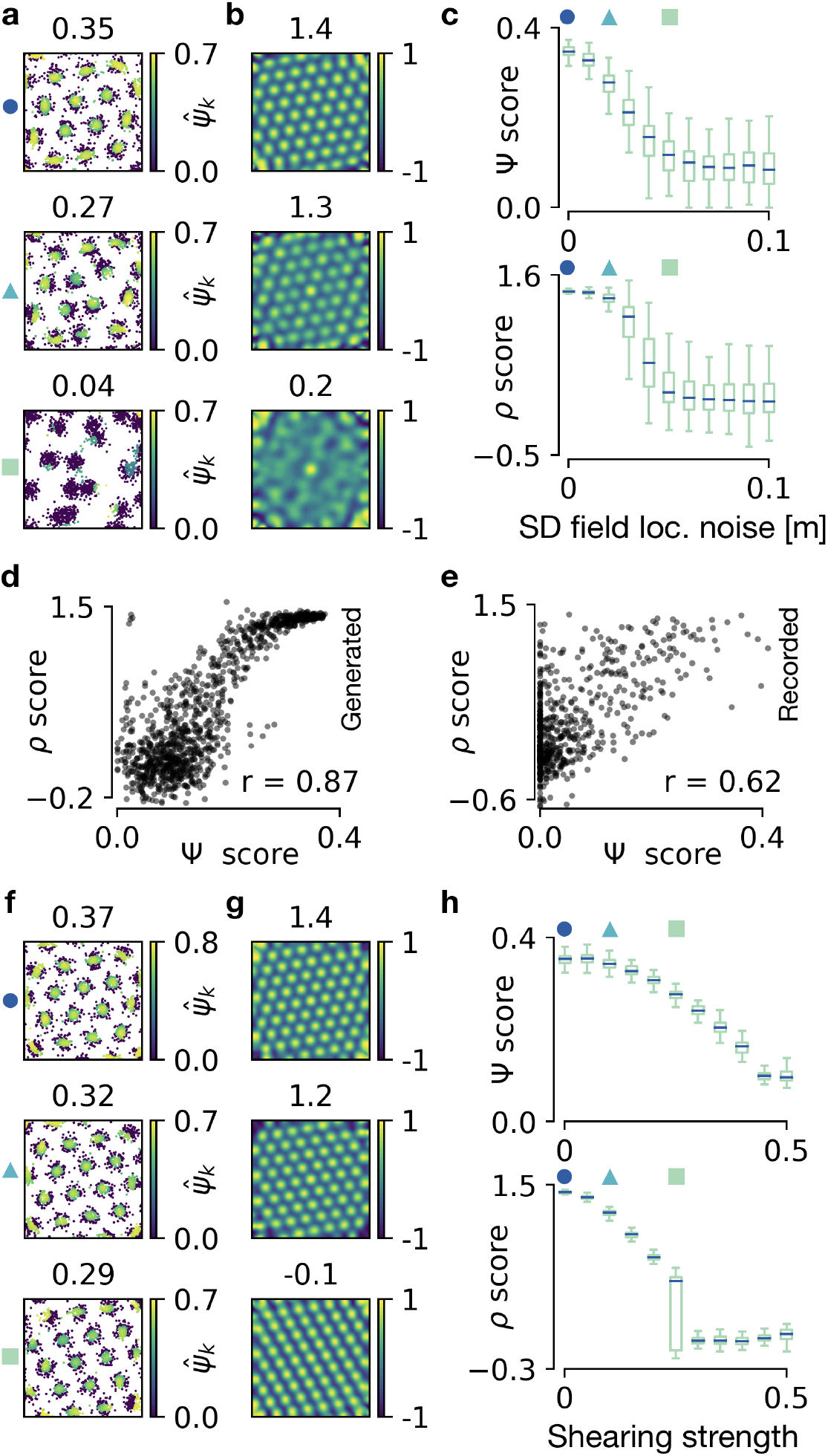
Quantifying the global symmetry of firing patterns. **a)** Generated spike maps in a 1m *×* 1m arena. Each spike k is color-coded with its grid score 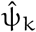; global ψ score shown above. Top: Perfect grid. Middle: Weakly perturbed grid field locations. Bottom: Strong perturbations. See text for description of perturbations. **b)** Autocorrelograms of the spike maps shown in a; ρ score shown above. **c)** Decay of ψ score (top) and ρ score (bottom) with increasing noise on grid field locations. Box plot for 100 random realizations at each noise level. For details on the box plot see Section S.4. Noise values for the examples shown in a are indicated with markers. **d)** Correlation of score with ρ score and Pearson’s coefficient r for the generated data used in a,b,c. **e)** Same as d but with experimentally recorded data from (Sargolini et al., 2006a,*b*). **f,g)** Arrangement as in a,b, but for generated spike maps with different degrees of shearing. Top: No shearing. Middle: Weak shearing. Bottom: Strong shearing. **h)** Arrangement as in c but with increasing shearing strength on the horizontal axis. Shearing strengths for the examples shown in f are indicated with markers.

The two scores differ in how they react to small and medium perturbations of the field locations. The median of ψ scores decreases linearly with increasing standard deviation of the grid field locations. In contrast, the median of ρ scores is not affected by small perturbations but decays abruptly at medium-sized perturbations. This non-linear decay arises because perturbations of field locations that are much smaller than the grid spacing lead to a blurred, but almost perfectly hexagonal correlogram (Figure 2b). Larger perturbations destroy the symmetry of the correlogram.

To study whether the two scores agree on the gridness of individual firing patterns, we correlate their values. The resulting Pearson correlation coefficients are r = 0.87 for generated data and r = 0.62 for experimentally recorded data (Figure 2d,e; data from Sargolini et al., 2006a,*b*). The lower correlation for experimental recordings results mainly from firing patterns for which either of the two scores failed in detecting the required shell (the neighborhood shell for the score and the annulus for the ρ score; see also Figure 3).

**Figure 3:**
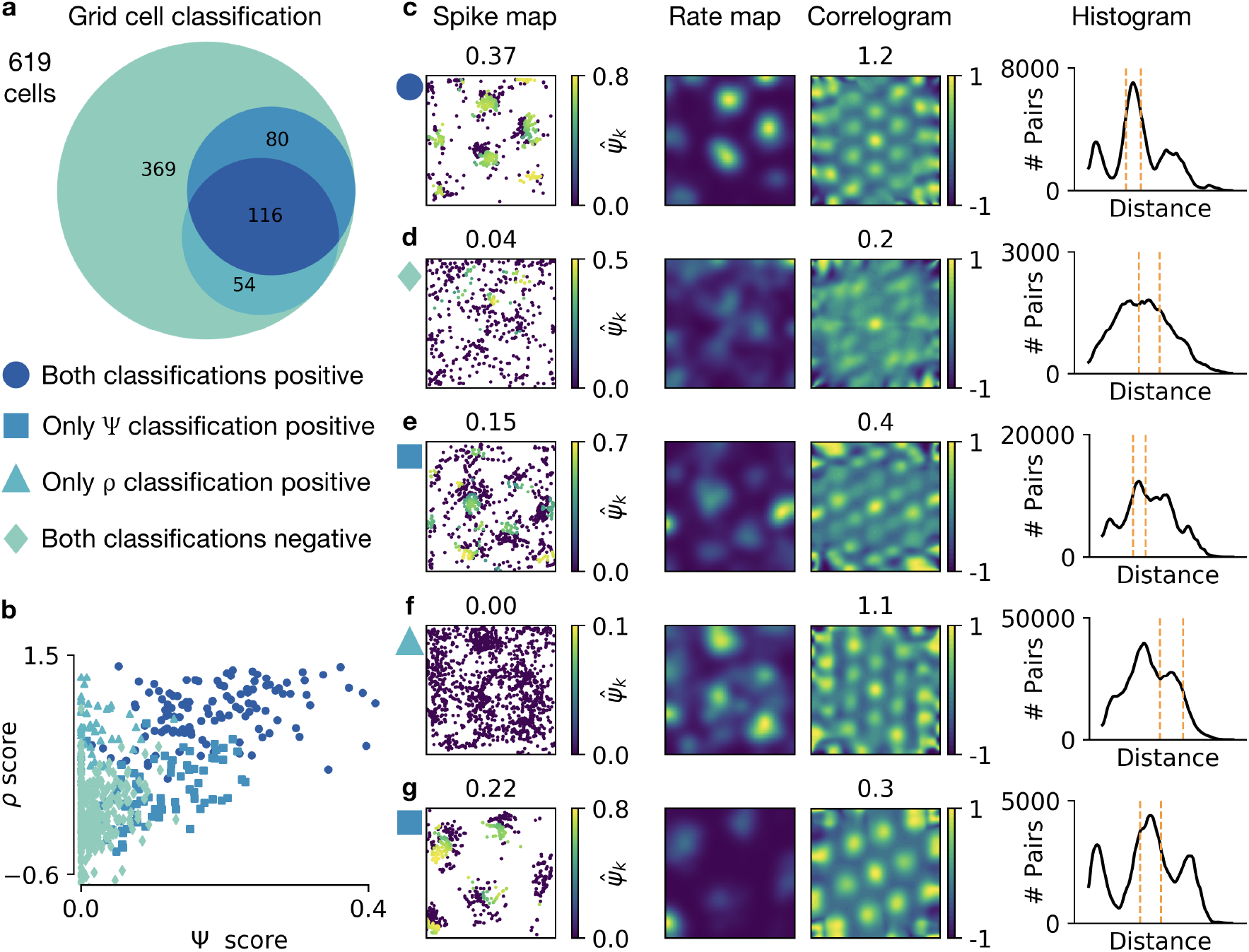
Classifying grid cells with the spike-based and the correlogram-based grid score. **a)** Venn diagram of 619 recordings from cells in mEC. Different colors highlight areas with different classification results (classification by ψ and/or ρ score positive/negative). See text for a description of the classification method. **b)** Correlation of ψ and ρ score as in Figure 2e. Markers represent the classification of the cell using the color code and the symbols from a. **c,d,e,f,g)** Individual recordings in different classification categories. The category is indicated by the blueish marker on the left, using the scheme in a. From left to right: spike map with each spike k color-coded with its grid score 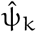 (global ψ score shown above); smoothed spike density (“rate map”), brighter values indicate higher density of spikes; autocorrelogram of the rate map (ρ score shown above); smoothed histogram of pairwise distances of all spikes, dashed orange lines show automatically detected neighborhood shells. See text for a detailed description of all examples. Experimental data recorded and made publicly available (Sargolini et al., 2006b) by Sargolini et al. (2006a).

Above, we distorted grid patterns by random shifts to each node of a hexagonal grid. Experiments have shown that grid patterns are often distorted in a different way: a perfect hexagonal grid is sheared and thus elliptically deformed (Stensola et al., 2015). To study the influence of elliptic distortions on ψ scores and ρ scores, we generate a perfect grid and apply a horizontal shearing transformation to the locations of grid fields (Figure 2, see Section S.2 for details). The shearing reduces the 6-fold symmetry and leads to a smooth and subtle decay of ψ scores with increasing shearing (Figure 2h). In contrast, the elliptic shape of the correlogram (Figure 2g) leads to a sudden drop of ρ scores (Figure 2h) at intermediate shearing strengths.

#### Classifying grid cells

When should we classify a recorded cell as a grid cell? A simple method for classification would be to call a cell a grid cell if its (ρ or ψ) score exceeds a fixed threshold. For the ρ score, a reasonable threshold is 0.75 (Kropff and Treves, 2008). From the relation between the two scores (Figure 2d,e), we could deduce a threshold of 0.15 for the ψ score. Using such a *fixed threshold* for classification can be misleading, though. A firing pattern with perfect hexagonal symmetry, but many “background” spikes outside of firing fields—which might just be an artifact of the extracellular recording, rather than a property of the cell—should be classified as a grid cell, even if its grid score is low. For grid cell classification it is thus common to use a *flexible threshold*, i.e., an individual threshold that is computed for each cell. It is determined as follows: the spike locations are shuffled at random (see Section S.1 for details on the shuffling procedure). This shuffling is done with 100 random initializations and results in 100 new spike maps. The grid score is computed for each shuffled spike map. The threshold is then defined as the 95th percentile of the grid scores obtained for the shuffled spike maps. Only if the grid score of the actual spike map exceeds this threshold, the cell is called a grid cell. This classification method can be used for either grid score.

For the experimental data from Sargolini et al. (2006a), which are publicly available (Sargolini et al., 2006b), we find that the ψ score and the ρ score lead to the same classification for more than 78% of the cells. Out of 619 cells, 196 are classified as grid cells by the ψ score and 170 by the ρ score (Figure 3a). 116 cells are classified as grid cells by both scores. Cells that are classified as grid cells by a score also tend to have a high value of this score, although sometimes even low grid scores result in a positive classification (Figure 3b).

For symmetric grid patterns with a small number of background spikes, the ψ score and the ρ score typically both classify the cell as a grid cell (Figure 3c). For non-grid cells with irregular firing, the classification of both scores is typically negative (Figure 3d).

A disagreement between the two scores can have different reasons. If the grid pattern has many spikes outside of grid fields, the cell might be classified as a grid cell by one or both scores although both are low (Figure 3e). For strong activity outside of firing fields, the detection of the correct neighborhood shell can fail, thus leading to wrongfully low grid scores of all spikes (Figure 3f). In line with the observed robustness of the ψ score to elliptic distortions (Figure 2h), cells with elongated grid patterns may be classified as grid cells by the ψ score, but not by the ρ score (Figure 3g).

In summary, the ψ score and the ρ score perform comparably in quantifying and classifying grid cells, but the ψ score is more robust to elliptic distortions. In the next section, we highlight the advantages of a spike-based grid score for the analysis of local grid properties.

### 3.2 The spike-based score detects local grid distortions

So far we assumed that gridness is an arena-wide property of a firing pattern. We did not consider local defects. But recent experiments have shown that grid patterns are often locally distorted. Such distortions have been observed in both asymmetric (Krupic, Bauza, Burton, Barry, et al., 2015; Krupic, Bauza, Burton, and O’Keefe, 2018) and quadratic environments (Hägglund et al., 2018; Stensola et al., 2015), and are presumably caused by boundary effects. Moreover, Wernle et al. (2018) have shown that grid patterns recorded in two arenas separated by a wall do not form a coherent grid (Wernle et al., 2018), but merge into a more coherent pattern when the wall is removed.

We now show that our spike-based grid score is suitable to quantify and visualize such local properties. We first illustrate this for generated spike data and then for experimental recordings.

To generate an example of a grid pattern with local defects, we first generated a perfect grid and then added random vectors to the field locations (cf. Figure 2a, see Section S.2 for details) only in the eastern half of the arena (Figure 4a,b). To analyze gridness locally, we divide the arena into partitions and compute the ψ score within each partition, i.e., the mean over all ψ̂ _k_ scores for all spikes k within the given partition (Figure 4c). Averaging over multiple cells highlights that the gridness decays along the west-east axis but stays stable along the south-north axis, and this trend is already visible for individual cells (Figure 4c,d).

**Figure 4:**
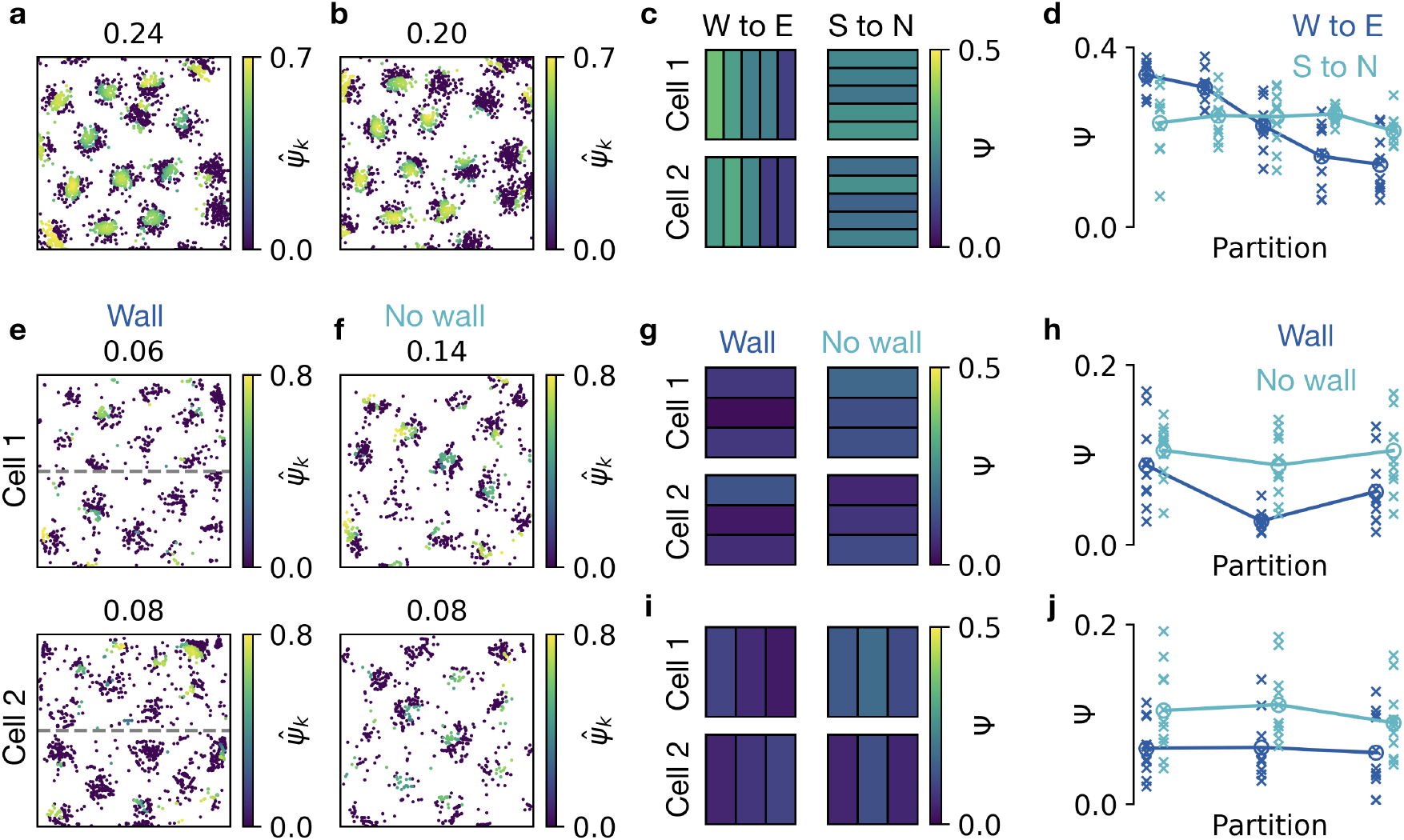
Visualizing and quantifying local distortions of grid patterns. **a,b)** Generated spike locations with noise on grid field locations applied only on the east side of the arena (see text for details, 2 example cells). The color of each spike location shows the 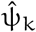 score. Shown above: ψ score. **c)** East-west (left) and south-north (right) partitioning of the arena. Black lines show partition boundaries. Colors of partitions indicate the score within the partition for the examples in a (top) and b (bottom). **d)** ψ score in partitions along west-east axis (dark blue) and south-north axis (light blue). Crosses show results for 10 randomly generated spike maps with noise on field locations applied only on the east side. Filled circles indicate the mean over 10 realizations. **e)** Spike locations of two example cells recorded in a rat that explored the northern and southern half of a box. The two halves are separated by a wall (dashed gray line). The spikes in each half were recorded in separate sessions. Color-coding as in a. **f)** Spike locations of the two cells shown in e while the rat explored the entire arena after the separating wall was removed. **g)** Grid score in south-north partitions before wall removal (left) and after wall removal (right) for the cells in e,f. Color-coding as in c. **h)** ψ scores in the south-north partitions shown in g for 11 recorded cells. Crosses denote individual cells, filled circles the mean across cells. ψ scores before wall removal in dark blue,Ψ scores after wall removal in light blue. **i)** ψ scores in west-east partitions for the cells in e,f. Arrangement as in g. **j)** ψ scores in the west-east partitions shown in **i**. Same recordings and arrangement as in h. Data in panels **e-j** kindly provided by (Wernle et al., 2018).

We now apply the same analysis to recent experimental data. Wernle et al. (2018) recorded grid cells from rats that explored the northern and southern half of a 2m *×* 2m box (Wernle et al., 2018). The two halves were initially separated by a wall. The cells fired in grid patterns independently in both halves of the arena (Figure 4e). Then the authors removed the wall and let the rat explore the entire arena. At the southern and northern boundaries, the grid fields typically stayed at the same locations as before wall removal. Close to the former location of the wall, the grid fields moved—and sometimes even merged—to form a more coherent grid pattern (Figure 4f). Quantifying the local increase of coherence requires a local grid score.

We therefore use our spike-based grid score to show that the rearrangement of grid fields after wall removal indeed leads to a local increase of gridness. Given that the firing patterns of the grid cells are noisy and strongly elliptic, it is difficult to determine the peaks in the histograms of pairwise spike distances (Figure 4e,f). We thus set the central distance of the neighborhood shell manually to ℓ = 65cm—a value that we obtained by visual inspection of the histograms of pairwise spike distances of all 11 recorded cells, after wall removal. We divide the arena into three equal sized partitions. For south-north partitioning (Figure 4g), the grid score is typically lowest in the partition that contains the separating wall (Figure 4h). After wall removal, the grid score increases in all partitions, but the increase is most prominent in the partition that contained the separating wall (Figure 4h). In contrast, a west-east partitioning does not lead to a single partition with predominantly low grid scores. Moreover, the increase after wall removal is similar for all west-east partitions (Figure 4i,j). Our analysis thus quantifies the qualitative observation that two initially independent grids locally rearrange to form a more coherent grid. For less elliptic grid patterns, we assume our quantitative results to be even more pronounced.

In summary, a spike-based grid score visually highlights local grid defects. Averaging spike scores in arbitrary partitions and potentially over many cells enables a systematic analysis of grid defects on a spatial scale smaller than the grid spacing.

### 3.3 The spike-based score detects shifts in grid orientation

Early work on grid cells has shown that the firing patterns of anatomically nearby cells typically have a similar orientation (Hafting et al., 2005), which tends to align with a cardinal axis of the environment (Krupic, Bauza, Burton, Barry, et al., 2015; Stensola et al., 2015). While these findings suggest a constant grid orientation, experiments from the same group have shown that the orientation can drift within the arena or exhibit local distortions (Hägglund et al., 2018; Stensola et al., 2015). To quantify these experiments, a local measure of grid orientation is needed.

In the following, we study local grid orientation using the angle θ_k_ of spike k (see Algorithm 1 for the definition) on generated (see Section S.2) and recorded firing patterns that were kindly provided by Stensola et al. (2015).

When each spike is color-coded according to its grid orientation θ_k_, the spikes of firing fields have the same color for a perfect hexagonal grid—except the spikes at the edge of a grid field (Figure 5a). In contrast, the color-coding shows a sudden switch if the orientation is changed abruptly (Figure 5b) or a color gradient if the grid orientation drifts continuously across the arena, both in generated data (Figure 5c) and in recordings (Figure 5d). If the drift in orientation is along the south-north axis of the box, the gradient follows this axis (Figure 5c,d). This can be highlighted by partitioning the arena and calculating the mean value of the orientation θ_k_ for all spikes k within each partition (Figure 5c,d). To highlight an orientation drift recorded in multiple cells, we average over the the orientation θ_k_ for spikes from several cells. For the recordings of Stensola et al. (2015), this shows that the orientation increases monotonically from south to north, but varies less systematically from west to east, for a set of 5 cells recorded in the same rat (Figure 5e,f). Note that changes in grid orientation distort the correlogram, which can lead to low ρ scores (Figure 5b,c), even if the grid-like structure is clearly apparent. In contrast, the ψ score is robust to distortions in orientation because of its locality.

**Figure 5:**
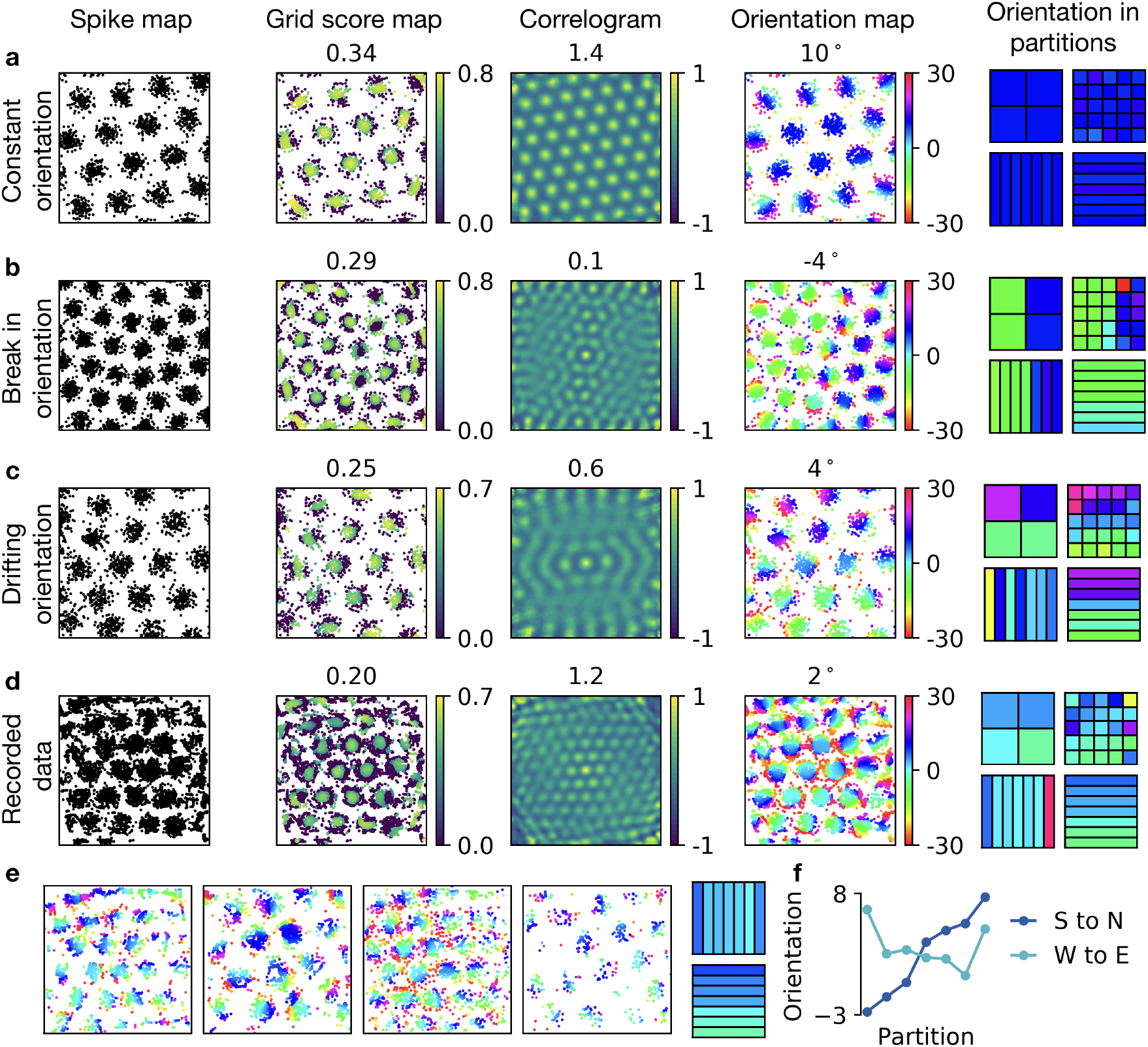
Visualizing and quantifying local changes in orientation of grid patterns. **a,b,c,d)** From left to right: spike locations; spike locations where each spike is color-coded with its 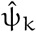 score (global ψ score shown above); autocorrelogram of the firing pattern (ρ score shown above); spike locations where each spike is color coded with its orientation θ_k_ (global orientation Θ shown above); four different spatial partitionings of the arena where black lines show partition boundaries and colors of partitions show the average orientation within each partition (mean of θ_k_ values) using the orientation color-code. **a)** Generated perfect grid pattern with orientation of 10 degrees. **b)** Generated grid pattern with abrupt change in orientation. **c)** Generated grid pattern with drifting orientation. **d)** Experimentally recorded grid pattern with drift in orientation. **e)** Four further experimental examples and the average of their spike-based orientation values in west-east and south-north partitionings. **f)** Orientations along west-east and south-north in the partitions shown in e. Data kindly provided by Stensola et al. (2015).

In summary, a spike-based orientation measure highlights local distortions in grid orientation. Averaging spike-based orientations, using arbitrary spatial partitionings, allows to further classify the distortion.

### 3.4 The spike-based score quantifies temporal fluctuations of grid patterns

Whether or not the spikes of a cell form a hexagonal spatial pattern can vary in time. For example, the activity of grid cells loses its periodicity when the light is turned off (Chen et al., 2016; Pérez-Escobar et al., 2016). A decay of grid-like firing patterns can also be triggered by internal modifications, like the deactivation of hippocampal drive to the entorhinal cortex (Bonnevie et al., 2013). In the following, we show how the spike-based grid score can be used to study changes in grid cell activity with high temporal resolution.

The standard procedure for computing a grid score in a given time interval is to take the spike locations from all spikes that occurred in this interval and compute the grid score for the resulting pattern. While this can be done with both the correlogram-based score and the spike-based score, it introduces constraints on the temporal resolution, because the temporal window must contain a sufficient coverage of the arena to obtain a meaningful score.

This problem can be circumvented with a spike-based grid score, because for a given spike map, each spike k has a grid score 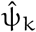 and a time t_k_. In other words, the grid score 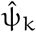 at time t_k_ quantifies the contribution of spike k to the grid pattern formed by *all* spikes. Just like the global grid score ψ is calculated by averaging over the individual 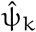 values of *all* spikes, a temporally local grid score can be obtained by averaging the 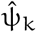 values of the spikes in an arbitrary time interval. The resulting average quantifies how much the spikes in this interval contribute to the grid in the complete recording.

To demonstrate that the spike-based grid score can characterize temporal changes in grid patterns, we consider generated data with a sudden transition from spatially homogeneous to grid cell activity (Figure 6a). Plotting the individual grid scores as a function of spike time highlights the transition to the grid pattern, although the grid scores 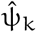 of the individual spikes are computed using the locations of all spikes (Figure 6a,b). The transition is emphasized by temporal smoothing (Figure 6c). Requirements for the smoothing filter are discussed later. Note that such an analysis first of all requires that a grid pattern can be detected in the full recording, so that a meaningful neighborhood shell can be identified.

**Figure 6:**
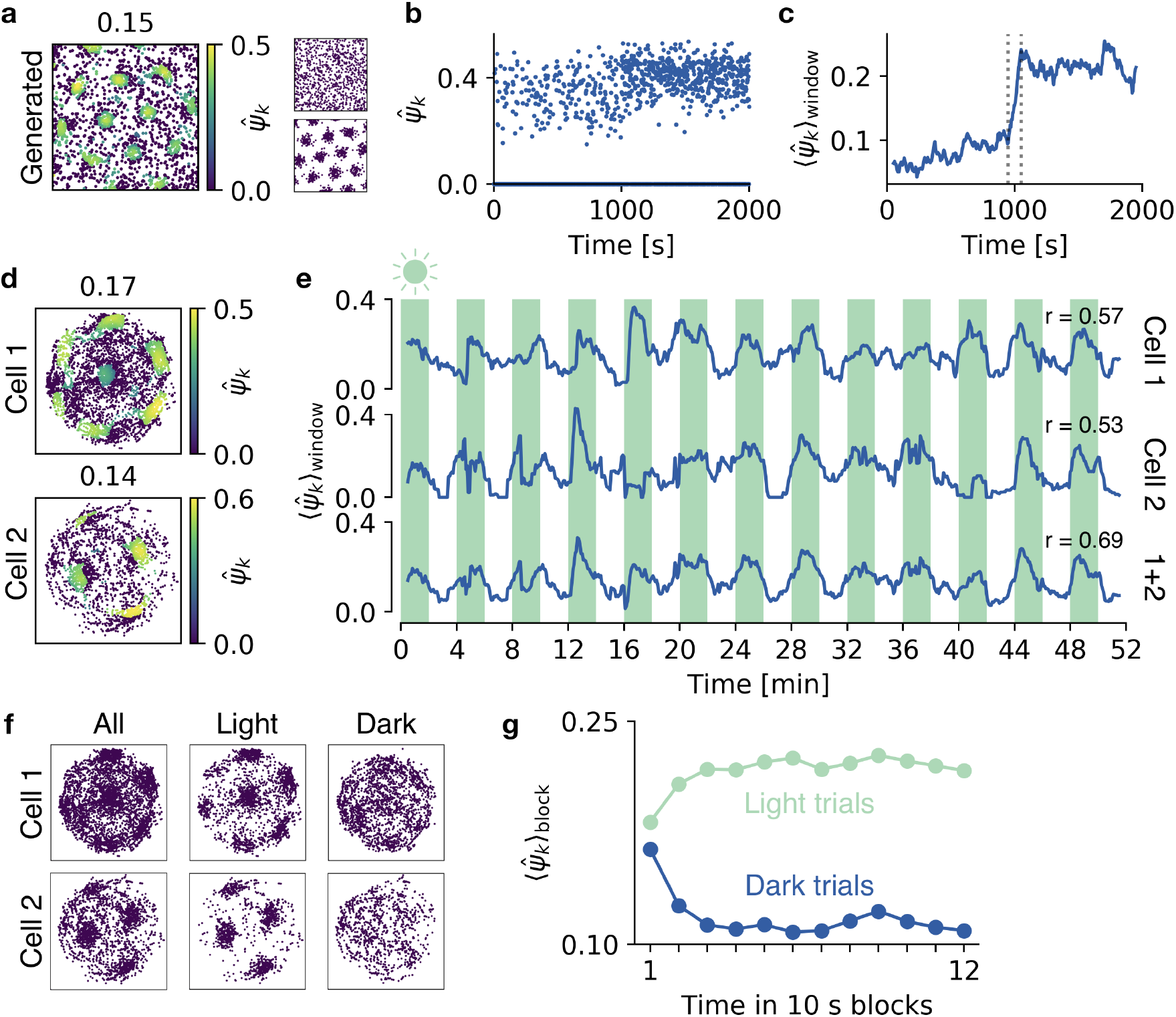
Quantifying temporal disruptions of grid patterns. **a)** Generated spike locations with 2000 spikes (left). Each spike k is color-coded with its grid score 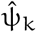. The first 1000 spikes are drawn from a spatially uniform distribution (top right) and the last 1000 spikes are drawn from a perfect grid (bottom right), at a constant rate of 1 spike per second. **b)** 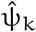 scores in a as a function of spike time. Note that the score of many spikes is 0. **c)** The scores from b, filtered with a 100 second time window (dotted lines indicate window size). **d)** Two experimentally recorded spike maps, color coded with 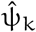 scores. The recordings are 52 minutes in total and contain 13 trials in light (with an illuminated cue card) and 13 trials in darkness, each of 2 minutes duration. **e)** The 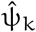 scores as a function of time, filtered with a 1 minute window. From top to bottom: First cell from d; second cell from d; mean over both cells. Shadings indicate light trials. Pearson’s correlation coefficients, r, between lighting condition and filtered grid scores are shown for each scenario. **f)** Spike maps for cell 1 and cell 2, separated into spikes fired during light trials and spikes fired during dark trials. **g)** Temporal evolution of grid scores after changes in lighting condition. Each 2 minute trial was separated into 12 *×* 10 second blocks. Each point shows the mean 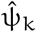 score of all spikes in a time block in light trials (light blue) and dark trials (dark blue), averaged over 73 grid cells. Grid scores are calculated in reference to all spikes during light trials (see text for details).

We now use this method to analyze spike patterns from cells in mice that forage in an arena in alternating trials of darkness and light (Pérez-Escobar et al., 2016). For the recorded grid cells (Figure 6d), the filtered time evolution of grid scores 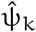 is strongly correlated with darkness and light—grid scores tend to be higher in light trials (Figure 6e). The Pearson correlation coefficient between lighting condition and the smoothed grid scores (r = 0.57 and r = 0.53 for single cells and r = 0.69 for the average over smoothed grid scores of two cells) indicate that only few simultaneously recorded cells are necessary to reach confident conclusions about temporal variations in the quality of grid patterns or contextual influences on grid cell activity.

But how quickly does the grid pattern decay when the light is turned off? How rapidly does it reappear when the light is turned on again? Pérez-Escobar et al. (2016) answered these questions with a temporal resolution of 10 seconds using *map similarity* Experimental data recorded and made publicly available by Pérez-Escobar et al. (2016). (Pérez-Escobar et al., 2016) (see below for a discussion of map similarity). We now show that we can reach the same temporal resolution using the spike-based grid score.

To this end, we create a *reference spike map* for each recorded grid cell in the data by Pérez-Escobar et al. (2016), which comprises the locations of all spikes that were fired during light trials. Note that these subsets of spikes show a much clearer grid pattern than that of spikes during darkness (Figure 6f). We then quantify how well the spikes of a given cell fit into the grid pattern of the cell’s reference spike map, by adding every spike k *individually* to the reference spike map and computing its 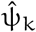 score as usual.

We then average the 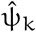 score of spikes that occur at similar times during light/dark trials. To this end, we divide each 2 minute trial into 12 *×* 10 second blocks. This division is done separately for light and dark trials. We then average the grid scores 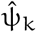 over all spikes and over 73 recorded grid cells within each 10 second block. Plotting this average as a function of time shows that grids decay within 30 seconds after the light is turned off (Figure 6g). Grids need a similar time to reappear after the light is turned on (Figure 6g).

In summary, the spike-based grid score readily contains temporal information which can be used to study distortions of the grid pattern in time. The time evolution of the grid score can be correlated with any other time varying variable to unravel its influence on the grid pattern, e.g., light conditions, running speed, concentration of neuromodulators.

## 4 DISCUSSION

In summary, we have developed a method to quantify the embedding of each spike of a cell into a hexagonal neighborhood of spikes. The method yields both a local grid score and a local grid orientation. It is based on the ψ^(M)^ measure for M-fold symmetry used in solid state physics (Halperin and Nelson, 1978) and adapted to the analysis of spike locations. It requires only few and robust steps to assign a grid score to each spike. We have shown that the global average of this spike-based score is as adequate as existing correlogram-based methods to quantify and classify grid cells. Its key advantage is that it simplifies local analyses both in space (‘Where are good spikes?’) and in time (‘When are good spikes?’). The method allows a new visualization of grid cell data, showing all spike locations together with information on local grid quality and local orientation.

### 4.1 Averaging over multiple cells increases spatial or temporal resolution

We have shown spatially local grid scores in sub-partitions of the arena and temporally local grid scores in time intervals. The resulting values can be misleading if the spatial subinterval is too small or if the temporal subinterval is too short. For example, if we study the local grid score of a single cell in a partition that does not contain a grid field of that cell, the mean over all spike scores in that partition will be very low, suggesting a local defect. Such sampling biases could be mitigated by making the partition larger (or the interval longer, for temporal analyses), at the cost of resolution. A reduction of sampling biases without a loss of resolution can be obtained by averaging over multiple cells. It would be beneficial if these cells have different spatial phases, because this ensures the presence of a grid field in each partition and thus reduces sampling biases. Averaging over many cells thus enables a higher spatial resolution than the grid spacing and a higher temporal resolution than the time the rat needs to sample a sufficient number of grid fields in a single cell.

### 4.2 Comparison with correlogram-based method in spatial subregions

The correlogram-based ρ score can be made more local by computing it in spatial subregions within the arena. A window can also be used to compute the sliding average of the ρ score in the entire arena (Hägglund et al., 2018), resulting in a spatial map of local grid scores. Selecting the size of the window is subject to a trade-off: while a small window is necessary for a high spatial resolution, the ρ score will fail to detect a grid if the window is too small—e.g., smaller then the grid spacing. In contrast, the spike-based score can be computed in arbitrary spatial partitionings (Figure 5) by averaging over the spikes in each partition. The side length of a partition can even be smaller than the grid spacing, in particular if multiple cells are used to compute the score in each partition (see above). The flexibility to use partitionings of an arbitrary shape, e.g., trapezoidal or triangular, can be important to study grid cells in environments with complex geometries (Krupic, Bauza, Burton, Barry, et al., 2015; Krupic, Bauza, Burton, and O’Keefe, 2018).

### 4.3 Comparison with map similarity for the study of time evolution

We have shown that a spike-based grid score enables a straightforward analysis of the temporal stability of grid patterns and illustrated this on recorded data from mice foraging through an arena in light and darkness (Pérez-Escobar et al., 2016). In our analysis of the decay and reappearance of grid cell activity (Figure 6), we reproduced the results shown in Pérez-Escobar et al. (2016), where the authors used *map similarity* for their analysis. In the map similarity measure, a rate map is correlated with a reference map. As a reference map, the authors used recordings during times of light. In each 10 second block, they collected all spike locations from that block in all trials (separating dark and light trials) and used these spike locations to create a rate map. If the correlation of the resulting rate map with the reference map is high, the pattern is similar, indicating that the grid has not decayed yet, and vice versa. For grid patterns without defects in field locations and with similar firing rates for different grid fields, map similarity and our measure should lead to similar results. However, our measure focuses on how much a spike is part of the *grid* of the reference map. In contrast, map similarity only quantifies how much spikes are part of the *rate pattern* of the reference map, irrespective of whether it forms a grid. If for example, the reference map has a firing field that does not belong to the hexagonal pattern, spikes that are located in this firing field will lead to low values in our measure. In contrast, map similarity assigns high values to activity within any firing field of the reference map, be it a grid field or a defect.

### 4.4 Detecting grids with elliptic distortions

We have shown that the spike-based ψ score is more robust to elliptic deformations of grid patterns than the correlogram-based ρ score (Figure 2h). This robustness can be crucial for the detection of grid cells, because many grid patterns are of elliptic shape. Quantifying the local improvement of a grid pattern in the scenario where two grids coalesce in a contiguous environment (Figure 4) was only possible because grid patterns could be detected, even though they were strongly elliptic.

A correlogram-based grid score can also be used with elliptic grid patterns, but requires an explicit compensation of the ellipticity. To this end, the ellipticity is detected and removed from the correlogram before the grid score is computed, by compressing the correlogram along the principal axis of the elliptic distortion (Brandon et al., 2011). In principle, we could apply such a transformation to all spike locations and then compute the spike-based grid scores. We have shown, however, that such an additional step is not necessary as long as the ellipticity is not too strong.

### 4.5 Non-constant grid spacing

If a firing pattern is grid-like, but with different grid spacings in different parts of the arena, we cannot compute the neighborhood shell from all pairwise distances. Instead, the neighborhood would need to be determined individually for each spike, using the histogram of pairwise distances only between a given spike and all other spikes. Since we are not aware of local distortions to the grid spacing, we did not consider this.

### 4.6 Non-isotropic assignment of grid scores to spikes in boundary grid fields

For grid fields with six neighboring fields, spikes in the center of the field receive the highest grid score and grid scores decay isotropically in all direction. In contrast, in grid fields at the boundaries, spikes with high 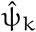 form a curved stripe (Figure S1a). The reason is the following: Being in the boundary grid field, moving towards the central neighbors perturbs the 6-fold symmetry stronger than moving tangentially to it (Figure S1b,c,d). Such a directional dependence is not present in grid fields that have neighboring grid fields on all sides.

### 4.7 The spike-based grid score for other symmetries

We focused on hexagonal patterns, i.e., grids of 6-fold symmetry. Experiments suggest that there are also band-like firing patterns, i.e., 2-fold symmetry (Krupic, Burgess, and O’Keefe, 2012)—though their existence has been questioned (Krupic, Burgess, and O’Keefe, 2015; Navratilova et al., 2016). Moreover, boundary effects in arenas of complex shape could cause grid deformations that result in other symmetries. The suggested spike-based grid score can be readily applied to any M-fold symmetry. For example, to study quadratic symmetry, we would replace M = 6 by M = 4 and compare to the remaining symmetries, now including 6 (see Algorithm 1). Depending on the symmetry, the radius of the neighborhood shell needs to be chosen differently. For example, for quadratic symmetry, results are improved if the neighborhood shell is set to 2/3ℓ instead of ℓ, to only incorporate the four nearest neighboring fields (Figure 7d). For band-like symmetry, the shell radius should be ℓ/2, so that the next band is not included in the shell (Figure 7e).

**Figure 7:**
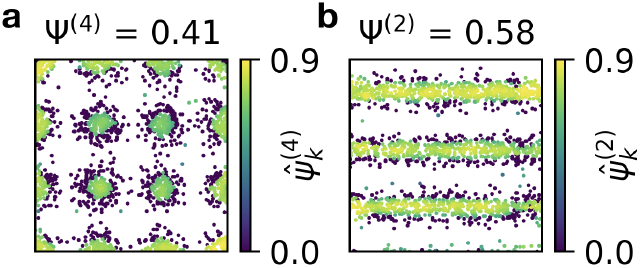
A spike-based grid score for other symmetries. **a)** Adaptation of the 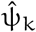 score to 4-fold symmetry. Shown above: average over all spikes. **b)** Adaptation of the 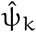 score to 2-fold symmetry. Shown above: average over all spikes. See text for details.

As the experiments on spatially tuned cells become more complex, new properties will be unveiled. A spike-based grid score could be used as a single unifying method to classify grid cells and to analyze their local and temporal characteristics. The presented method could further be useful for the analysis of grid patterns in other modalities (Aronov et al., 2017; Constantinescu et al., 2016) or in other contexts, e.g., in the anatomical arrangement of grid cells (Ray et al., 2014) or retinal ganglion cells (Jang and Paik, 2017).

## ACKNOWLEDGMENTS

We would like to thank Loreen Hertäg for proofreading this manuscript, Sophie Rosay for helpful discussions on correlogram-based grid scores and local grid defects, Martin Hägglund for helpful discussions on local grid defects and Tanja Wernle, Hanne and Tor Stensola and the Moser lab for generously providing their data.

Funding: Bundesministerium für Bildung und Forschung. Grant reference number: FKZ01GQ1201.

## SUPPORTING INFORMATION

### S.1 Shuffing spike locations for grid cell classification

When classifying grid cells, we use standard shuffling procedures that have been established for conventional correlation-based grid scores (Langston et al., 2010; Wills et al., 2010). To this end, shuffled spike locations are generated from the rat trajectory and the actual spike locations: The times of all spikes are shifted consistently by a temporal offset. This temporal offset is drawn randomly from the interval [20 seconds, time of last spike – 20 seconds] for 100 different shufflings. If, after shuffling, a spike time exceeds the time of the last spike in the original recording, its time is periodically wrapped to the beginning. The shuffled location of each spike is given by the position of the rat at the shifted time of the spike. This shuffling procedure assigns a new location to each spike while maintaining the temporal structure. We classify a cell as a grid cell according to the ψ score if the ψ score of the original recording exceeds the 95th percentile of all Ψ scores computed for the 100 shufflings.

**Figure S1:**
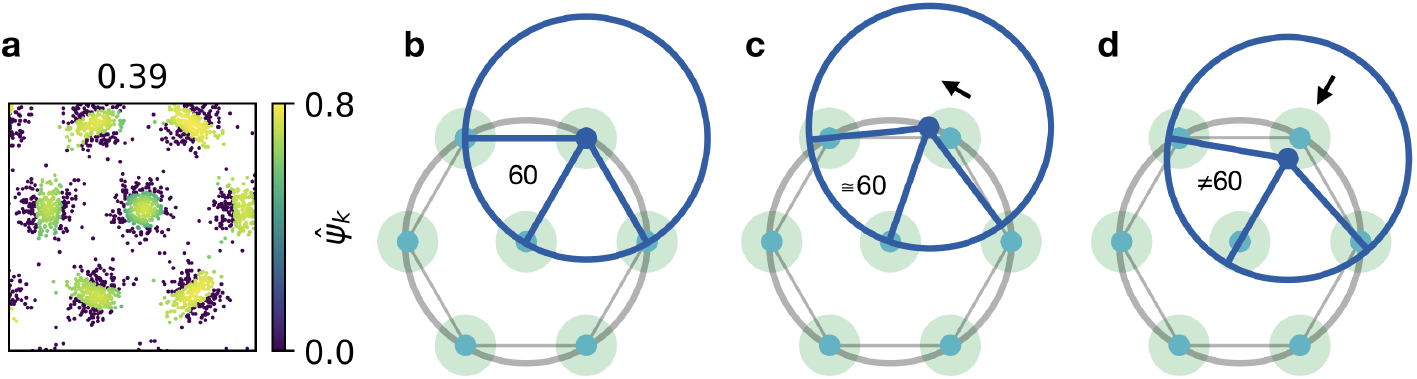
Explanation of non-isotropic assignment of grid scores to spikes in boundary grid fields. **a)** Generated spike locations with few firing fields. The color of each spike k indicates the 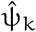 score. Fields at the boundaries have only three neighboring fields. This leads to a stripe-like assignment of 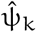 scores. **b)** Schematic of a perfect grid cell. Dots on the hexagon mark the centers of grid fields. Shaded regions show the grid field size. The dark blue dot in the upper right marks a reference spike in the center of a grid field. The dark blue circle around it indicates the neighborhood shell—much thinner than actual, for easier visualization. Connecting the reference spike to the center of the cross section of the neighborhood shell and the neighboring grid fields forms angles of 60 degrees. **c)** Same arrangement as in b but now the reference spike is not in the center of the grid field, but shifted along the gray circle that circumscribes the hexagon (direction indicated by arrow). Connecting the reference spike to the center of the cross section of the neighborhood shell and the neighboring grid fields still forms angles of roughly 60 degrees. Consequently, spikes along this direction have high 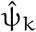 scores. **d)** Same arrangement as in b,c but now the reference spike is shifted orthogonally to the circle that circumscribes the hexagon. Connecting the reference spike to the center of the cross section of the neighborhood shell and the neighboring grid fields does not form angles of 60 degrees. Consequently, spikes along this direction have low 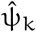 scores. For central grid fields, this effect is compensated by grid fields on opposite sides.

Correspondingly, to classify a cell as a grid cell according to the ρ score, we use the same procedure but with the ρ score instead of the ψ score. Note that, recently, it was suggested to use a different shuffling procedure that corresponds to randomly relocating the grid fields (Barry and Burgess, 2017).

### S.2 Generating artificial spike locations

To generate artificial grid patterns, we draw spikes from a probability distribution that is comprised of two-dimensional Gaussians whose centers lie on a—possibly perturbed—hexagonal grid. The center location of each Gaussian is the location of a grid field. For all generated data, we use an arena size of 1m *×* 1m. To comply with the experimental observation that the size of grid fields scales with the grid spacing (Brun et al., 2008), we use a constant ratio of the width of the Gaussians and the grid spacing before adding perturbations:

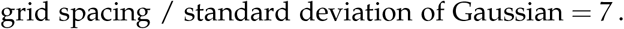

To assess the robustness of the grid score, we study several perturbations of hexagonal firing patterns. To this end, we first generate grid field locations on a perfect hexagonal grid. To avoid edge effects, this grid is generated on a much larger area that contains a perfect hexagonal arrangement of 271 (i.e., side length 10) grid fields, and the actual arena size (1m *×* 1m) is cut out in the center after the perturbations are applied.

In the scenario of random perturbations (Figure 2a), we add a different random vector to each grid field location. Each such two dimensional vector is drawn from a two dimensional normal distribution. The standard deviation of this distribution dictates the noise on the field locations (Figure 2c).

In the scenario of shearing, we apply a horizontal shearing transformation to each field location of the perfect lattice, by multiplying the two dimensional location of each firing field with the matrix

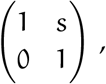

where s is the strength of the shearing (Figure 2h).

In the scenario of a drift in orientation along the south-north axis, we rotate the location of the i-th grid field (x_i_, y_i_)^T^ via

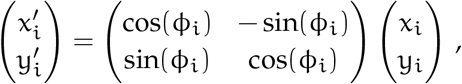

where 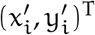 denotes the location of the grid field after rotation and ϕ_i_ increases along the south-north axis in the following way:

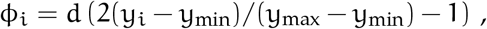

where y_min_ and y_max_ are the y-locations of the southernmost and northernmost grid fields, respectively. The factor d determines the strength of the drift. We use d = π/3, in the example shown in Figure 5c.

Finally, in the scenario of a sudden change in orientation (Figure 5b), we extracted the field locations from simulations by Rosay et al. (2018).

### S.3 The correlogram-based grid score

To compute the correlogram-based grid score, we use the method suggested by Langston et al. (2010). To this end, we first calculate a “rate map” by summing over Gaussian kernels with a width of 5% of the arena side length, centered at the locations of the spikes. We then determine the spatial autocorrelogram, i.e., the Pearson correlation coefficients for all spatial shifts of the rate map against itself. From this correlogram, we crop out an annulus that contains the six fields that are arranged around the central peak. To get the inner radius of this annulus, we clip all values in the correlogram that are smaller than 0.1 to 0. We obtain the resulting clusters of values that are larger than 0.1 using `scipy.ndimage.measurements.label` from the *SciPy* package for *Python* with a quadratic filter structure, ((1, 1, 1), (1, 1, 1), (1, 1, 1)), for a correlogram with 201 *×* 201 pixels. We use the distance from the center to the outermost pixel of the innermost cluster as the inner radius of the annulus. We obtain the outer radius of the annulus by trying 50 values, linearly increasing from the inner radius to a corner of the arena. With each of the resulting 50 annuli, we crop the correlogram, rotate it and correlate it with an unrotated copy of the correlogram. We determine the correlation coefficient for 30, 60, 90, 120 and 150 degrees. We define the grid score as the minimum of the correlation values at 60 and 120 degrees minus the maximum of the correlation values at 30, 90 and 150 degrees. After trying all 50 annuli, we take the highest resulting grid score as the grid score of the cell. A hexagonal symmetry thus leads to positive values whereas a quadratic symmetry leads to negative values. Since the correlogram-based score uses Pearson correlation values that are typically expressed with the Greek letter ρ, we refer to it as the ρ score. Note that the match of the grid score symbols with the first names of the authors that introduced them (ρ-samund and ψ-mon) is coincidental.

### S.4 Box plot settings

In all shown box plots, each box extends from the first to the third quartile, with a dark blue line at the median. The lower whisker reaches from the lowest data point still within 1.5 IQR of the lower quartile, and the upper whisker reaches to the highest data point still within 1.5 IQR of the upper quartile, where IQR is the inter quartile range between the third and first quartile.

